# BioMARathons as a seasonal engagement model for marine citizen science: adapting BioBlitzes to challenging coastal environments

**DOI:** 10.64898/2026.05.13.724939

**Authors:** Sonia Liñán, Berta Companys, Ana Álvarez, Meritxell Turó, Carlos Rodero, Xavier Salvador, Jaume Piera

**Affiliations:** EMBIMOS Research Group, Institute of Marine Sciences (ICM-CSIC), Spanish National Research Council (CSIC), Barcelona, Spain

**Keywords:** Marine citizen science, BioBlitz, citizen engagement, biodiversity monitoring, community reactivation, Janus Engagement Framework

## Abstract

BioBlitzes are widely used citizen science events that combine biodiversity monitoring, public participation, and environmental awareness through short and intensive observation campaigns. However, applying this model to marine environments presents additional challenges related to safety, access, weather dependency, specialised equipment, species identification, and sustained participation. This paper presents the BioMARathon model as a case study of how BioBlitz-inspired events can be adapted to marine citizen science contexts.

The BioMARathon extends the conventional BioBlitz format into a longer, seasonal, and distributed engagement model designed specifically for marine and coastal environments. The paper describes the conceptual foundations of the model in the Janus Engagement Framework, which informed both the design of the BioMARathon and the adaptation of the MINKA citizen science observatory to better support participation, validation, feedback, and continuity over time. BioMARató Catalunya, launched in 2021, is presented as the founding implementation of the model and as the basis for later replication in Portugal.

Between 2021 and 2025, BioMARató Catalunya showed continued growth in participation, observations, and taxonomic coverage, while also contributing to the detection of non-indigenous species, first regional records, and climate-related ecological impacts. Beyond biodiversity outcomes, the case suggests that extending participation across a season, distributing activities through local mobilising organisations, and combining expert validation with visible feedback mechanisms can support recurrent participation, retention, and community reactivation in marine citizen science. Rather than offering a formal causal evaluation, this article contributes practical lessons for the design of citizen science initiatives in challenging environments.

## Introduction

BioBlitzes have become a powerful tool in citizen science. BioBlitzes are collaborative events that bring together scientists, naturalists and members of the public to catalogue local biodiversity in an intense short period of time, usually from 24 hours to a few days (Roger and Klistorner, 2016). These events aim to record the maximum number of living species within the designated area and time. Since the success of the first BioBlitz in 1996 at Kenilworth Aquatic Gardens (Postles and Bartlett, 2018), many organisations around the world have replicated this participatory, short-duration event concept (Tiago, Evaristo and Pinto, 2024).

One of the key advantages of a BioBlitz is the generation of large amounts of data in a short period of time. Despite the initial idea that they could be a lot of work with limited scientific gain, BioBlitzes have increasingly been shown to provide valuable results from both academic and social perspectives (Aristeidou *et al*., 2021; Meeus *et al*., 2023; Tiago, Evaristo and Pinto, 2024). BioBlitzes, for example, can allow the finding of underrecorded species in the target area (Silvertown, 2009; Ballard, Dixon and Harris, 2016). Systematic analysis of BioBlitzes has shown that these events can involve over 120 participants that generate more than 2.000 observations from 300 species (Meeus *et al*., 2023).

Beyond data collection, BioBlitzes are effective events for science education and public outreach. After being involved, BioBlitzes participants report that they have increased their biodiversity and ecological knowledge, as well as developed a greater appreciation for local biodiversity (Ballard, Dixon and Harris, 2016). Complementing this, BioBlitzes foster hands-on learning by enabling participants to learn new identification species skills and put into practice the scientific method (Bonney *et al*., 2016). Moreover, BioBlitzes can enhance community cohesion and stewardship, as seen by the rise in volunteers and sustained monitoring initiatives in communities after participating in these events (Johnson *et al*., 2014).

In addition to data collection and education, BioBlitzes have shown their potential to increase public support for conservation through fostering direct connections between people and their local environments (Cooper, Shirk and Zuckerberg, 2014). The collaborative nature of these activities helps to build trust between scientists and the public, and encourages continued engagement in biodiversity monitoring and environmental conservation.

However, organising a BioBlitz is complex, and it requires considerable effort to fulfil all the logistics. A successful BioBlitz relies on the engagement of staff and participants in the planning and organisation of the events. These logistics tasks often include identifying and securing a location, obtaining the legal permissions, implementing safety measures, etc. Also, an effective communication team is needed for designing and executing a dissemination strategy to enhance community engagement and involvement in the event. Depending on the geographic scale, it may be appropriate to seek additional resources through sponsorship.

In conclusion, BioBlitzes show the successful integration of science and community involvement in improving biodiversity knowledge and community stewardship.

### A global reference for BioBlitzes: the City Nature Challenge

The City Nature Challenge (CNC) is one of the most popular global IT-based BioBlitzes, with thousands of individuals engaged in urban biodiversity monitoring. We understand as IT-based BioBlitzes those that use technological and digital platforms to run the event. Thanks to the IT services (managing citizens’ participation and data quality control), it is possible to engage thousands of participants and also provide fast methods for data quality control.

CNC was initially planned in 2016 as a friendly competition between San Francisco and Los Angeles, but has grown swiftly, with over 400 cities in over 40 countries competing annually (Palma *et al*., 2024). The event gives communities a framework to record local biodiversity and invites the public to get involved in conservation initiatives.

Why is it an IT-based BioBlitz? Because the CNC enables participants to upload observations to citizen science observatories such as iNaturalist, MINKA or any other. A global community of specialists and enthusiasts validate the observations. In particular, the CNC has been useful for local governments as it provides an affordable method to assess urban biodiversity, guide land management decisions, and promote environmental stewardship at the community level (Palma *et al*., 2024).

CNC has also been educationally impactful, as participants have been able to have hands-on learning experiences to help them understand local ecosystems. Comparative research in many countries has demonstrated that CNC enhances the sense of connectedness with nature, particularly in urban people, where direct contact with biodiversity is often limited (Sakurai *et al*., 2022). Participants from different cultures have become more conscious of environmental issues and more motivated to take part in conservation efforts after their engagement in CNC (Sakurai *et al*., 2022). CNC engagement remained resilient even during the COVID-19 epidemic and community science approaches were adapted efficiently (Kishimoto and Kobori, 2021).

### Challenges and opportunities for marine citizen science

Marine citizen science (MCS) is gaining prominence in marine and coastal management (Cigliano *et al*., 2015), spatial ecology and conservation (Bosso *et al*., 2024). Participatory marine monitoring systems enable volunteers, from amateur naturalists to trained divers, to report data and expand their scientific knowledge through biodiversity monitoring, marine debris tracking, and environmental parameter collection (Busch *et al*., 2016; Ceccaroni *et al*., 2020; Kelly *et al*., 2020; Garcia-Soto *et al*., 2021; Soacha Godoy *et al*., 2022). These participatory activities enable data gathering in previously understudied marine areas and enhanced public involvement in marine conservation issues (Thiel *et al*., 2014; Dean *et al*., 2018).

Marine citizen science initiatives have shown significant policy significance and research potential. For example, citizen-produced data from the UK’s *Seasearch* project helped to designate 38 marine conservation zones (Earp and Liconti, 2020). Similarly, the Marine Biodiversity and Climate Change project (MarClim) has produced data that is important evidence of changes in species distributions owing to climate change, and offers practical guidance for adaptive marine management and policy makers (Soacha Godoy *et al*., 2022). These examples highlight the promise of MCS to impact marine governance and to empower coastal populations to participate in ocean stewardship at the same time.

MCS is under-represented in the literature compared to terrestrial projects (Roy *et al*., 2012; Sandahl and Tøttrup, 2020; Garcia-Soto *et al*., 2021; Wehn *et al*., 2025). But the quick growth of this field highlights its increasing importance to the academic and the general public. Volunteer contributions provide rich coverage of maritime ecosystems, which may be difficult and/or costly to access using traditional scientific methods and represent a major opportunity to complement professional research efforts (Kelly *et al*., 2020). In addition, MCS facilitates the co-creation of ecological knowledge, aligning with the agendas of inclusive and transformative research (Fraisl *et al*., 2020; Soacha Godoy *et al*., 2022).

However, MCS has its own unique obstacles, largely due to the intrinsic complexity and logistics of marine environments compared with terrestrial ones. Furthermore, participation is limited by logistical issues, such as the requirement for certain equipment (e.g. fins, diving masks, wetsuits or underwater cameras) or safety issues (Martin *et al*., 2016). This, together with the fact that MCS sometimes needs technical skills such as swimming or diving, as well as access to intertidal and underwater environments, can influence the motivation and long-term engagement of participants (Martin *et al*., 2016; Liñán *et al*., 2022). Similarly, social or geographical constraints might influence the inclusiveness and representativeness of participation, which requires enhancing community outreach and support mechanisms (Mazumdar *et al*., 2018).

The nature of marine species and their habitats makes it more challenging to engage the public in marine biodiversity monitoring. Non-specialists generally don’t know many taxonomic groups of marine organisms. Even if they are phylogenetically distant, MCS volunteers may have difficulty in distinguishing between coralline algae, bryozoans and sponges (Cigliano *et al*., 2015; Terenzini, Safaya and Falkenberg, 2023). In terrestrial settings, on the other hand, unskilled and non-expert participants may discriminate between obvious classifications such as birds and reptiles. This information gap and the limited access to underwater environments limit the opportunities for broad participation in MCS, unlike land-based efforts.

Unlike the terrestrial initiatives, where the participants can typically post their observations to a citizen science platform digitally on site, the underwater data collection usually requires the participants to get out of the water to re-establish the connectivity before uploading their findings. This additional process prevents immediacy and can decrease motivation, resulting in data loss (Terenzini, Safaya and Falkenberg, 2023). Additionally, maritime habitats lack obvious terrestrial markers, and the absence of distinct site boundaries can impede participants from developing a sense of ownership and attachment to a certain spot, which may result in reduced involvement (Cigliano *et al*., 2015).

The unpredictability of weather and sea conditions adds an additional dimension of risk to MCS events. Activities may be cancelled and data continuity and engagement affected by adverse sea conditions, such as strong currents or poor underwater visibility.

MCS also faces the challenge of maintaining the quality and consistency of data gathered by volunteers. Inconsistencies in methods of observation, data input procedures and species identification accuracy might undermine the reliability of results, especially when protocols are not standardised (Ceccaroni *et al*., 2020; Kelly *et al*., 2020). Standardised procedures and accurate data validation mechanisms, which are missing in many present programs, are essential for the proper incorporation of MCS into national or international biodiversity databases.

MCS involves logistical and safety challenges that aren’t as visible in terrestrial citizen science. Scuba diving, snorkelling, boat-based activities and specialised instruments increase operational costs and risks, requiring strict safety regulations and insurance considerations (Cigliano *et al*., 2015). These factors tend to concentrate participation among skilled ocean users such as divers, recreational boaters or marine biologists and may reduce participation from wider audiences (e.g. tourists).

MCS integration into legislative and regulatory frameworks is still at its infancy. Successful case studies have demonstrated the impact of volunteer data on marine management. However, many citizen science datasets remain underutilised, due to concerns about their quality or their incompatibility with formal monitoring programs (Fraisl *et al*., 2020; Gumiero *et al*., 2025). This needs building partnerships between scientists, citizen science networks, local authorities and policymakers to ensure that participatory data are validated and contextualised within decision-making processes.

To address these MSC problems for both volunteers and organisers, a variety of techniques is required. Effective outreach and communication strategies are especially important to boost volunteer motivation and engagement. Transparency about the participation process, the participants’ role, and the concrete impacts of their contributions influences the feeling of purpose and ownership of the participants (Liñán *et al*., 2022). Ongoing communication through social media, community events, and dedicated engagement teams stresses the value of volunteer involvement and promotes sustained engagement and a sense of community among volunteers (Cigliano *et al*., 2015; Mazumdar *et al*., 2018).

## The BioMARathon Model: Adapting BioBlitzes to Marine Citizen Science

To address some of the specific constraints of MSC, the BioMARathon model adapts the short, intensive logic of the BioBlitz to a longer, seasonal and distributed format. While “Blitz” evokes a sudden collective effort to reach a specific goal, the term “BioMARathon” combines the idea of a longer-lasting process with “MAR”, meaning sea in Catalan, to emphasise the marine focus of the initiative.

The BioMARathon is an international citizen science event designed to document marine and coastal biodiversity through local, regional or national editions organised within a shared framework. Participants upload biodiversity observations to a citizen science platform, while local organisations, scientific institutions and public authorities support mobilisation, validation, curation, learning and dissemination. Its purpose is not only to record species, but also to create the conditions to foster recurrent participation, marine biodiversity learning and the production of biodiversity data that can support research, awareness-raising and conservation.

But the BioMARathon goes beyond an extended BioBlitz; it is a new class of citizen science event resting on four interrelated design principles (Figure 1). First, participation was extended across a season rather than concentrated into a single short event. Second, activity was distributed across multiple local opportunities for involvement rather than being tied to a single site and date. Third, the initiative relied on a visible network of local mobilising partners rather than on centralised recruitment alone. Fourth, participation was supported by relatively rapid scientific feedback and validation, helping volunteers see that their contributions were part of a meaningful collective process.

**Figure 1:**
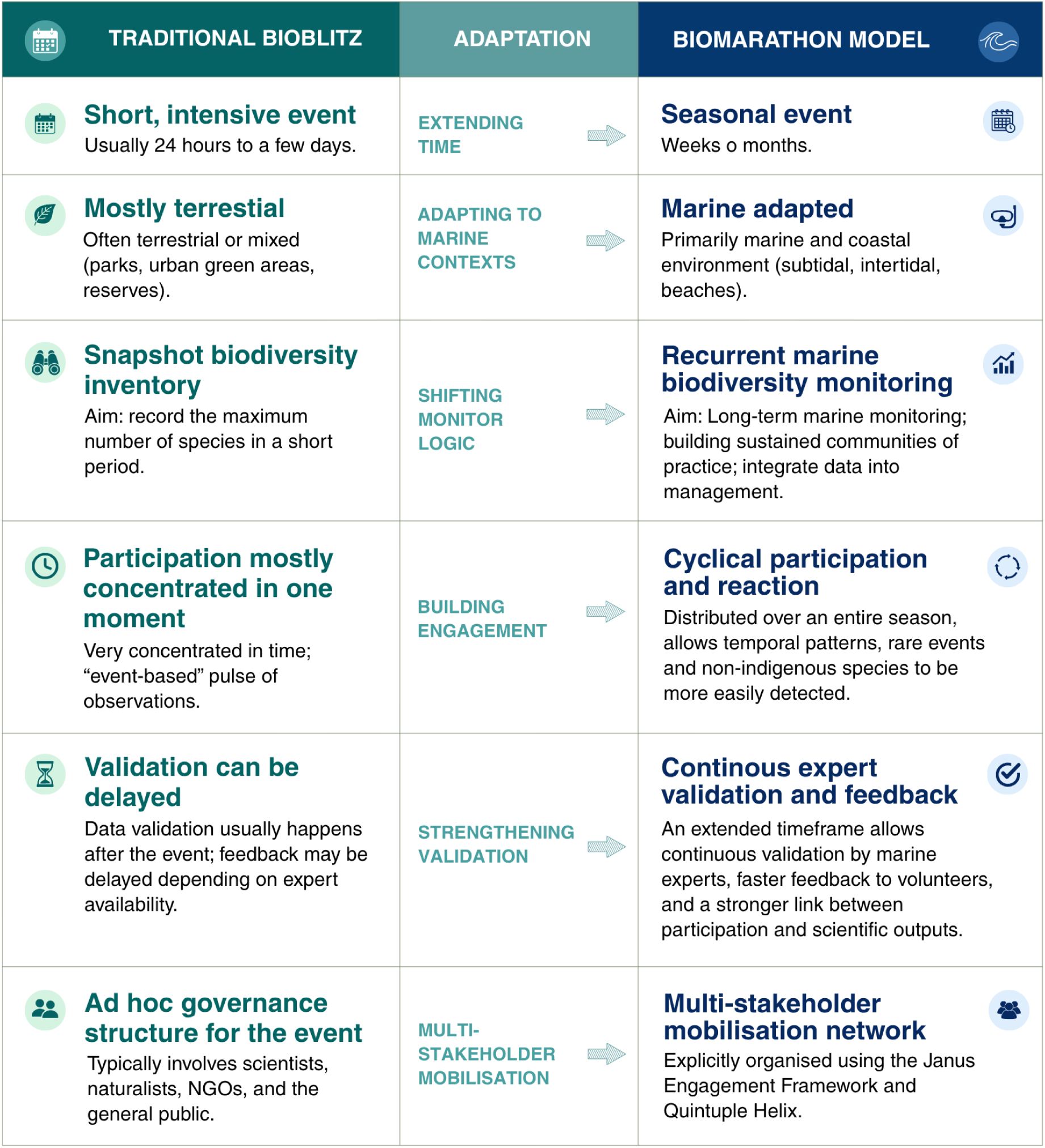
Comparison between BioBlitz and BioMARathons.

The first design choice mattered because marine participation is highly sensitive to environmental conditions. By spreading the initiative across months instead of days, BioMARathon reduced its dependence on a single weather window and increased the chances that different kinds of participants could join at least some of the activities. This extension also aligned the event more closely with ecological seasonality: a longer campaign provides an opportunity to register a wider range of marine organisms observations and seasonal variation over a brief snapshot could reasonably provide.

The second and third design choices mattered for engagement. A short event can be intense and motivating, but it also produces a very narrow window for participation. BioMARathons instead created multiple points of entry through guided outings, local communication, and repeated opportunities to contribute. This is especially important for marine contexts, where many potential participants depend on trusted intermediaries such as diving centres, federations, or coastal organisations in order to access the sea safely and meaningfully. The mobilising community, therefore, did more than advertise the initiative; it acted as a bridge between scientific aims and real participation conditions.

Rapid feedback also plays a central role. The original project design identified fast validation or identification responses as one of the key factors supporting volunteer engagement (Liñán *et al*., 2022). That is relevant in marine citizen science because the field setting itself often prevents immediate upload and recognition. If participants only receive delayed or opaque responses, the perceived value of their contribution can weaken. The extended seasonal design of BioMARathons gave experts more room to validate observations continuously, making feedback part of the engagement logic rather than an afterthought.

Also, a longer data collection time increases the likelihood of recording a wider range of species, including non-indigenous species, protected species, and rare occurrences that may not be observed in a brief monitoring period. As a result, the findings communicated to participants are more comprehensive and scientifically valuable, offering richer insights into marine biodiversity. This increases both the robustness of the data and the participant experience, as the more extensive and impressive results are a powerful motivator for continued engagement.

### The Janus Engagement Framework

To implement the BioMARathon, we use the Janus Engagement Framework (Figure 2), originally developed to address long-term public engagement challenges in citizen science projects (Liñán *et al*., 2022). In the BioMARathon model, the value of Janus lies in its capacity to organise engagement as a distributed process involving different communities, responsibilities and forms of motivation. This is especially relevant in MCS, where barriers to participation are not only motivational but also logistical, technical, social and environmental.

**Figure 2:**
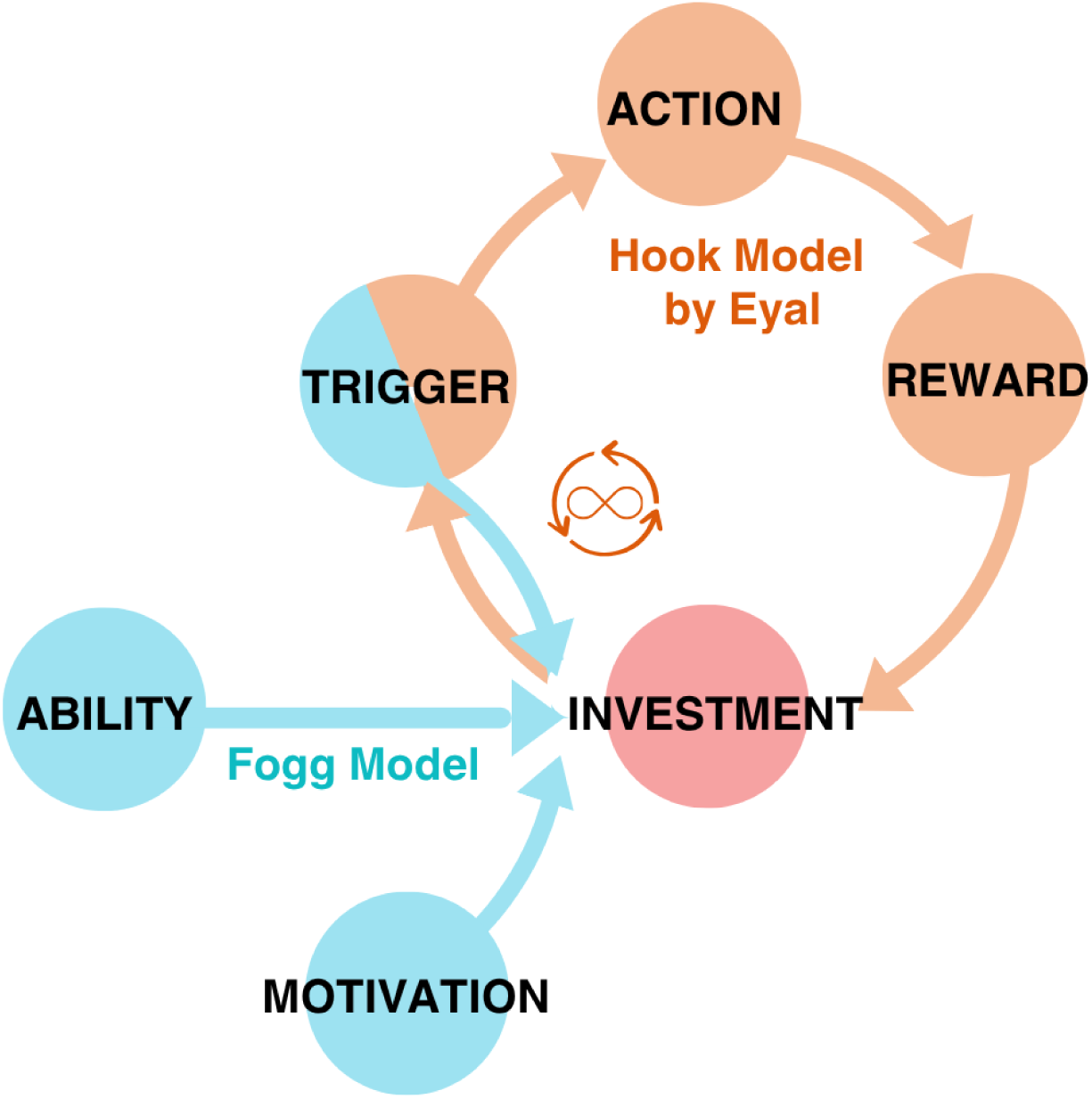
Simplified representation of the Janus Engagement Framework applied to the BioMARathon model. This figure illustrates the Janus Engagement Framework as the integration of the Hook Model and the Fogg Model. The Hook cycle links trigger, action, reward, and investment, while the Fogg Model highlights motivation and ability as conditions shaping participation. In the BioMARathon context, investment is central because it links initial engagement to future re-engagement, supporting continuity and retention across successive cycles of participation.

MCS initiatives must address several simultaneous challenges: access to the sea, weather and safety constraints, the need for specific equipment or skills, difficulties in identifying marine species, delayed data upload after underwater activities, and the need for trusted validation mechanisms. It is difficult for a single stakeholder to tackle all these issues. For this reason, BioMARathons built this engagement strategy on the Janus Framework, being designed as a multi-stakeholder engagement system in which different communities perform complementary roles. The Janus Framework helps to identify who can trigger participation, who can reduce barriers, who can provide learning and scientific credibility, and who can offer recognition or rewards that make sustained participation meaningful.

The framework applies the Quintuple Helix of Innovation (Carayannis, Barth and Campbell, 2012) to identify these roles and define four community types:

**The Participatory Community** includes the volunteer participants who contribute to the initiative as observers, identifiers, validators or local ambassadors. In BioMARathons, this community is diverse: it may include underwater photographers, divers, snorkellers, coastal naturalists, families, students and people interested in marine biodiversity. Their contribution is not limited to uploading observations; they also help to expand collective knowledge by learning to recognise species, improving data quality over time, sharing the initiative through their networks, and returning in subsequent editions. For this community, engagement is supported through clear participation instructions, accessible activities, visible results, species identification support, friendly competition, public recognition and the possibility of seeing their observations become part of a wider scientific and conservation effort.

**The Academic Community** includes universities, research centres and scientific experts responsible for the scientific framing of the initiative. In BioMARathons, this community provides methodological guidance, taxonomic expertise, data validation and curation, training and feedback. Academic actors, therefore, act both as scientific guarantors and as engagement enablers: they transform volunteer observations into validated biodiversity data, provide workshops and identification support, and create feedback loops that show participants that their contributions have scientific value. Rapid validation or identification responses, ideally within a short period after the observation is uploaded, are an important engagement mechanism because they convert participation into learning and recognition (Liñán *et al*., 2022).

**The Mobilising Community** includes the organisations that are in direct contact with potential participants and can activate participation locally. In BioMARathons, this community may include underwater activities federations, diving centres and clubs, snorkelling companies, environmental NGOs, local associations, schools, museums or coastal outreach organisations. Their role is central because many potential participants cannot access marine environments safely or confidently without trusted intermediaries. Mobilising actors organise guided activities, communicate the initiative to their own communities, help participants use the digital platform, provide local knowledge and reduce practical barriers such as access, equipment, safety and confidence. They also benefit from the initiative by offering their members or clients opportunities to collaborate with science, receive training from experts and participate in a collective environmental action.

**The Facilitating Community** includes public authorities, protected area managers, municipalities and other institutions responsible for the governance or stewardship of the study area. Their role is to create enabling conditions for participation and to connect the initiative with local environmental management, education or conservation agendas. In BioMARathons, facilitating actors can support permits, dissemination, access to public spaces, local legitimacy, institutional visibility and the potential use of citizen-generated data in decision-making. Their involvement can also reinforce the perceived relevance of the initiative, helping volunteers understand that their observations are not isolated records but part of a broader effort to improve knowledge and stewardship of marine and coastal ecosystems.

Figure 3 shows how this multi-stakeholder approach structured the BioMARathon as a dynamic Community of Practice (CoP), in which each stakeholder group played a distinct and necessary role; Figure 3 also complements this by illustrating concrete examples of the engagement actions through which these roles were operationalised in practice.

**Figure 3.**
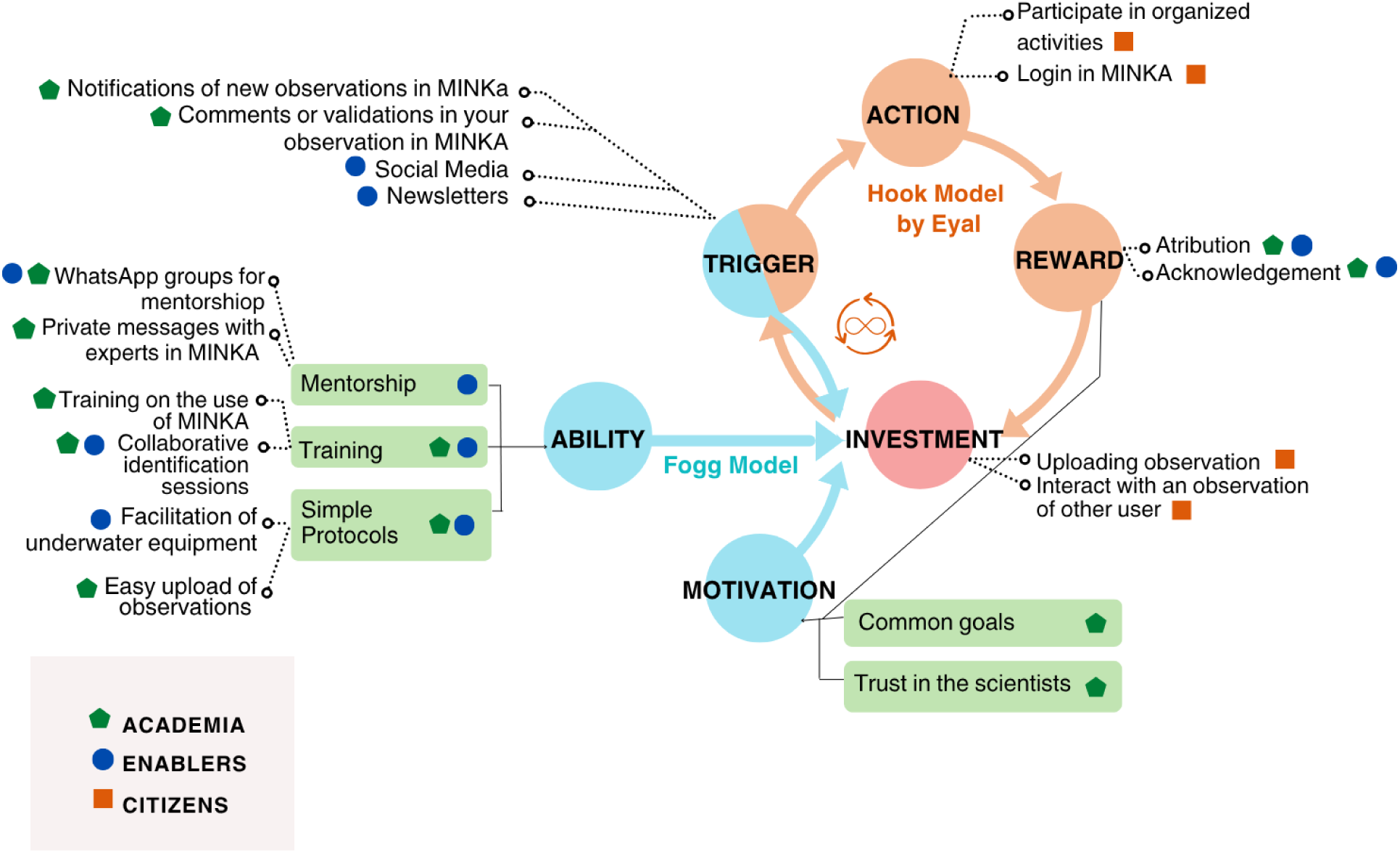
Examples of actions through which the Janus Engagement Framework was implemented in the BioMARathon and the MINKA observatory. This annotated figure presents concrete examples of actions associated with different elements of the Janus Engagement Framework in the BioMARathon, including notifications, comments and validations in MINKA, social media, newsletters, organised activities, log-in and observation upload, attribution and acknowledgement, mentorship, training, collaborative identification sessions, simple participation protocols, shared goals, and trust in scientists. The symbols indicate the stakeholder groups mainly involved in each action: academia, citizens, and enablers, where enablers correspond to the mobilising and facilitating communities in the Janus framework. The figure illustrates how engagement was operationalised through multiple coordinated actions across both the participatory process and the supporting observatory infrastructure.

### MINKA: a citizen science observatory for BioMARathons

MINKA (EMBIMOS research group, 2021) is the participatory infrastructure used to implement the BioMARathon concept. It was originally based on the iNaturalist code, which has become a reference for IT-based biodiversity recording and BioBlitz-style participation. However, in the BioMARathon model, MINKA is not only a data repository or an observation upload tool, but one of the infrastructures that makes the Janus Engagement Framework operational.

MINKA supports BioMARathons by enabling the different Janus communities to perform their roles within a shared digital environment. For the participatory community, the platform provides a visible space where volunteers can upload observations, receive identifications, follow the progress of the campaign and see their contributions as part of a collective biodiversity effort. For the academic community, MINKA provides tools to explore, filter, identify and validate observations, making it possible to transform volunteer contributions into more reliable biodiversity data. For the mobilising community, it offers a common platform with specific projects for their guided activities, training sessions and local campaigns. For the facilitating community, it provides a public and traceable record of biodiversity observations that can support awareness-raising, environmental education and, where appropriate, local management discussions. In this sense, MINKA complements the Janus Engagement Framework by providing the technical layer required for engagement actions to become visible, coordinated and sustained.

MINKA has also been adapted to better support marine biodiversity monitoring during BioMARathons (Figure 4). As an example, its graphical tools and filter panels allow observations to be selected according to predefined criteria. MINKA has incorporated dedicated filter options for marine taxonomic groups, facilitating the work of marine specialists who need to explore, select and validate different taxa during BioMARathons according to their expertise. This is particularly important in MCS because expert validation is both a data quality mechanism and an engagement mechanism: it improves the reliability of observations while also providing participants with feedback that reinforces learning and motivation.

**Figure 4.**
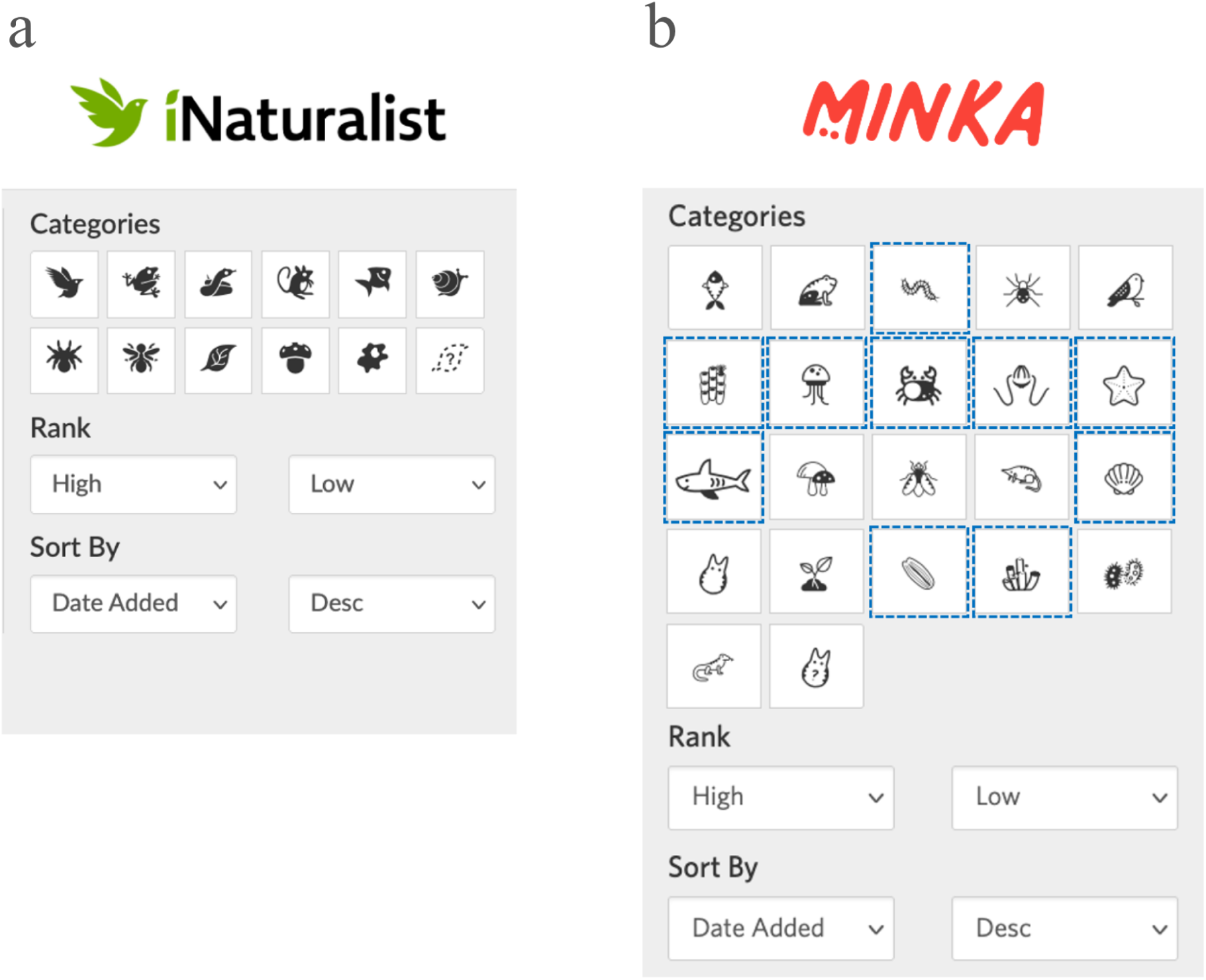
Example of an adaptation for managing marine observations in MINKA. (a) Initial filter panel for searching observations in iNaturalist, which is more focused on terrestrial taxons. (b) MINKA filter panel with specific filters for marine taxons (icons highlighted in dotted blue).

## BioMARató Catalunya: the Founding Case of the BioMARathon Model

The BioMARathon model is conceived as an umbrella format that can be implemented through different local, regional or national editions. In this sense, it follows a logic similar to other large-scale citizen science events such as the City Nature Challenge: a shared participatory structure, common digital infrastructure and comparable engagement principles are adapted to different territories through local organising communities. The first BioMARathon was launched in Catalonia in 2021 under the name BioMARató Catalunya. The model was later replicated in northern Portugal in 2024 as BioMARatona Norte and expanded into the whole Portuguese continental coast in the 2025 edition. Further expansion has been discussed in other regions.

BioMARató Catalunya is the founding implementation of this model. The first edition took place between spring and summer 2021 and involved the provinces of Barcelona, Girona and Tarragona (Catalonia, Spain) through snorkelling, scuba diving and coastal observation guided activities and also spontaneous participation not linked to any activity. Its objective is to photograph as many living organisms as possible along the Catalan coast, including birds, coastal plants and underwater species, and to upload these observations to the citizen science observatory MINKA. The initiative is promoted by the Institute of Marine Sciences (ICM-CSIC).

Since then, BioMARató Catalunya has shown continuous growth in participation, observations and taxonomic coverage. In 2021, 117 people participated and collected more than 10,000 observations of 1,061 species. In 2022, the Catalan edition involved 127 participants and generated more than 21,700 observations of 1,300 species. In 2023, BioMARató Catalunya exceeded 60,000 observations, tripling the previous year’s total, with contributions from more than 300 volunteers and 1,440 species recorded during that edition. In 2024, the initiative reached more than 91,000 research-grade observations and over 1,720 species, with the participation of 481 people. In 2025, the fifth edition further expanded the model: more than 520 people contributed over 94,000 observations to MINKA, and 2,040 species were identified. With these contributions, BioMARató Catalunya accumulated more than 380,000 observations and 2,870 documented species.

These results show that BioMARató Catalunya is not only an engagement event, but also a mechanism for producing relevant biodiversity knowledge. Several findings are particularly significant. The 2021 edition detected 24 non-indigenous species (NIS) along the Catalan coastline and recorded two first records in Catalonia. The 2022 edition registered four first records for the Catalan coast and also documented mortality events in red coral and gorgonian populations, consistent with concerns about the ecological effects of high temperatures and marine heatwaves. During the 2023 edition, participants detected 34 NIS and 40 protected or threatened species. The three editions together had censused around 1,900 marine species. The 2025 edition achieved the first complete spatial coverage of the Catalan coastline, in 10x10 km² grid cells resolution.

The four Janus communities contributed to BioMARató Catalunya’s success. The participatory community includes volunteers contributing observations and identifications. The academic community is centred on the Institute of Marine Science (ICM-CSIC), which supports scientific framing, species validation and data curation, and the broader design of the initiative. The mobilising community includes the Catalan Federation of Underwater Activities (FECDAS) and a set of small coastal companies and organisations promoting volunteer participation locally through guided outings, communication, and logistical support. The facilitating community included municipalities and public authorities with responsibilities connected to coastal territories and environmental awareness, such as the Area Metropolitana de Barcelona (AMB).

Figure 5 shows a clear seasonal rhythm. Observations increase during the BioMARató period, when sea conditions are usually more favourable, organised activities are concentrated, and communication efforts are stronger. These changes occur during winter, when sea temperature decreases, weather conditions are less favourable, and overall outdoor activity makes the marine environment less accessible. This seasonal decline should not be interpreted as a failure of engagement. In marine citizen science, a reduction in activity during less favourable months is expected.

**Figure 5:**
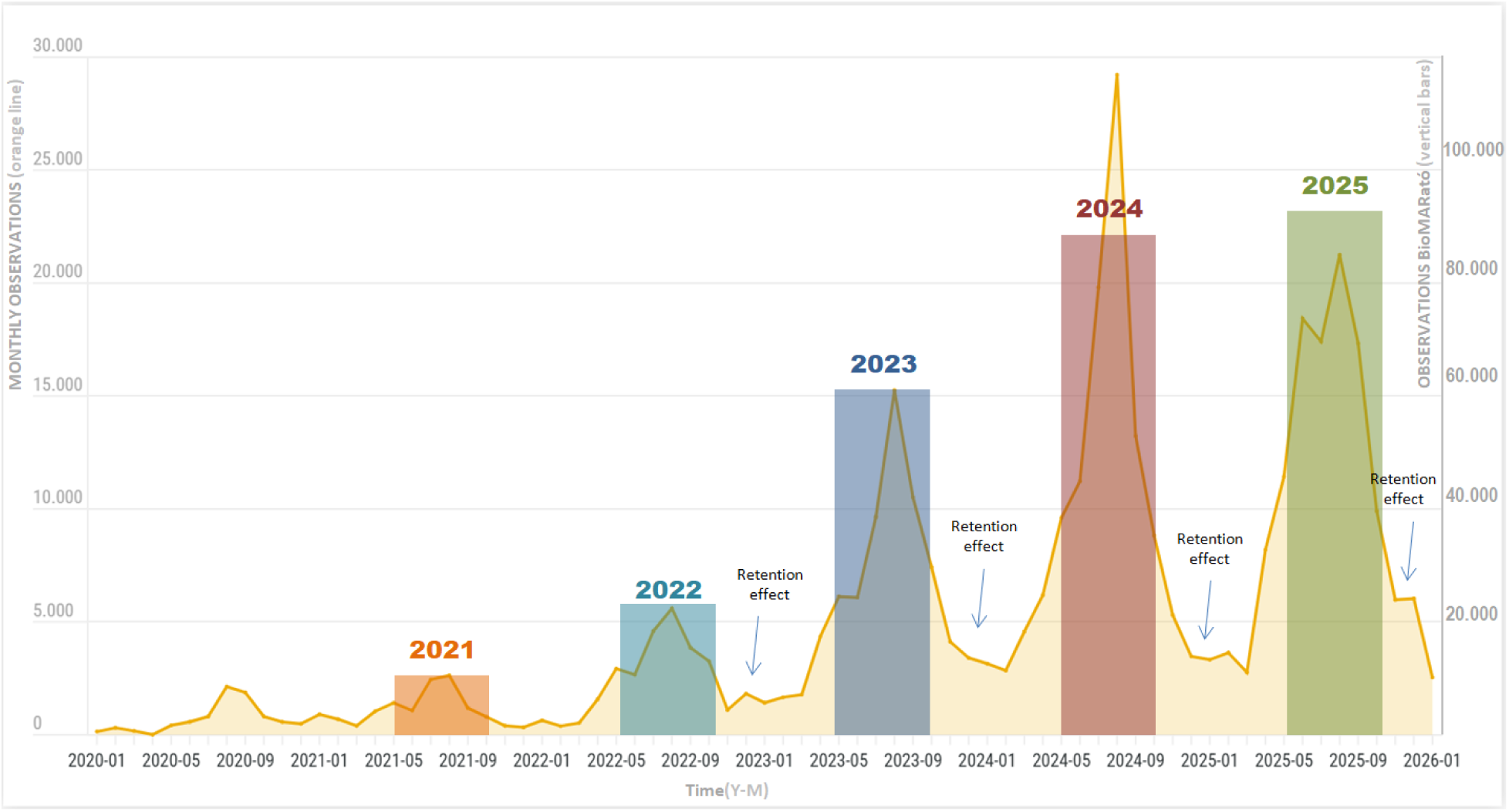
Evolution of the number of observations per month from January 2020 to January 2026 performed in the local BioMARathons (Catalonia coasts). The shaded vertical bands indicate the annual BioMARató periods, running approximately from the first Saturday of May until mid October. The figure shows the strong seasonal pattern of marine participation, with observation peaks during the BioMARató months and lower activity during winter. Rather than representing a single isolated annual event, the pattern suggests a cyclical but cumulative model of participation, in which each edition contributes to reactivating and consolidating the community of observers.

The key interpretive question is whether participation disappears after each edition or whether part of the community remains available to be reactivated. Figure 5 shows that after each BioMARató edition, activity does not simply return to the initial baseline observed before the initiative was consolidated. Instead, the figure suggests a cumulative pattern in which annual peaks are followed by lower but persistent activity, suggesting a retention rate of participants, and subsequent editions reactivate participation from a progressively stronger base.

This interpretation is important because the success of a recurring MCS initiative cannot be assessed only by the intensity of its annual peak. A short-term increase in observations during a campaign may indicate successful mobilisation, but it does not necessarily demonstrate sustained engagement. For BioMARató Catalunya, the relevant pattern is the combination of recurrence and accumulation: each edition acts as a new trigger for participation, while the persistence of activity between editions suggests that the initiative has gradually built a more stable community of marine observers.

This pattern also connects with the Janus Engagement Framework. Janus and its implementation combine short-term engagement mechanisms,such as campaign launches, guided activities, friendly competition, communication and public recognition, with longer-term mechanisms, including expert validation, learning, repeated participation, community identity and the integration of observations into a shared biodiversity dataset. In Janus terms, the annual event works as a recurring trigger, but retention depends on whether participants continue to perceive value after the immediate event has passed. Data suggests that BioMARató not only generate temporary mobilisation; it also creates conditions for reactivation, learning and continuity.

The 2025 edition is particularly relevant for interpreting this resilience. During July 2025, poor weather and bad sea conditions along the Catalan coast led to the cancellation of several BioMARató activities for safety reasons. In a short BioBlitz-style event, similar conditions could have seriously compromised the entire data collection window. In contrast, the extended BioMARathon format reduced dependence on a single favourable date or weekend. Although weather-related cancellations affected activity during part of the season, the overall pattern still shows continuity and reactivation. This supports one of the central arguments of the BioMARathon model: in marine environments, where participation is strongly conditioned by safety, access and environmental variability, engagement must be designed with enough temporal and organisational flexibility to absorb disruption.

## Lessons for Citizen Science Practice

BioMARató Catalunya suggests several lessons for designing citizen science initiatives in marine and other logistically demanding environments.

First, extending the duration of an event can be an engagement strategy, not only a sampling strategy. In marine contexts, participation is shaped by sea state, weather, accessibility, safety and seasonality. A longer event window reduces dependence on a single date and allows activities to be rescheduled when conditions are not safe or favourable.

Second, distributed local mobilisation is essential. BioMARató Catalunya shows that marine citizen science cannot rely only on centralised communication by a scientific institution. Mobilising organisations such as diving centres, federations, environmental NGOs and local associations act as trusted intermediaries. They provide access to participants, local knowledge, logistical support, safety conditions and social legitimacy.

Third, feedback and validation should be treated as engagement mechanisms. Expert validation is often discussed in relation to data quality, but in the BioMARathon model, it also supports learning, recognition and motivation. This is especially relevant in marine biodiversity, where many taxa are difficult for non-specialists to identify.

Fourth, digital infrastructure must support communities, not only data collection. MINKA enables volunteers, experts, mobilising organisations and facilitating institutions to act within a shared environment. It therefore functions as part of the engagement architecture, connecting observations, validation, learning and public visibility.

Finally, recurring citizen science events should consider retention and reactivation as key indicators of success. In seasonal marine contexts, sustained engagement does not necessarily mean constant participation throughout the year. It may appear as cyclical engagement: participants reduce activity when conditions are unfavourable but return during subsequent editions.

## Constraints and Limitations

The Catalan coast case study describes the development and implementation of the BioMARathon model rather than presenting a formal evaluation of its causal effects. The annual figures on participants, observations and species are descriptive indicators of growth and consolidation, but they should not be interpreted as experimental evidence.

The interpretation of retention is also based on community-level observation patterns rather than on a formal user-level retention analysis. Monthly observation data suggest reactivation and cumulative participation across editions, but future work should analyse individual participant trajectories to distinguish new participants, returning participants, occasional contributors and highly active long-term contributors.

The BioMARató Catalunya case is also shaped by specific territorial and institutional conditions, including the role of ICM-CSIC, the availability of MINKA, collaboration with diving and environmental organisations, and the characteristics of the Catalan coast. These conditions may not be directly replicable in all coastal regions. The BioMARathon should therefore be understood as an adaptable model rather than a fixed protocol.

Finally, the model requires sustained coordination. Extending an event over several months reduces dependence on a single weather window, but it also requires continuous communication, partner coordination, data validation and curation and platform maintenance.

## Conclusion

BioMARató Catalunya illustrates how a marine biodiversity event can be redesigned from a short BioBlitz-style activity into a seasonal BioMARathon model. Its value lies not only in extending the time available for observation, but in changing the engagement conditions around marine citizen science.

The BioMARathon model combines a longer event window, locally distributed mobilisation, expert validation, public recognition, and the support of the MINKA digital infrastructure to create different entry points for participation and encourage continued engagement over time. The case suggests that, in challenging environments where participation is shaped by weather, safety, access and technical skills, success should not be assessed only through short-term peaks of activity. The capacity to sustain connection and reactivate communities across editions may be equally important.

The BioMARathon model, therefore, offers a practical approach for designing citizen science events that are better aligned with the social and ecological realities of marine biodiversity monitoring.

## Author contributions

SL, Conceptualisation, Writing–original draft. BC, Data curation, Writing–review and editing. AA, Data analysis. XS, Data curation. MT, Writing–review and editing, CR, Writing–review and editing, JP: Conceptualisation, Writing–original draft, Funding acquisition.

## Funding

The EMBIMOS research group was funded by the European Commission through the PHAROS project, funded by the European Union’s Horizon Europe programme under Grant Agreement No. 101157936, the ANERIS project under Grant Agreement No. 101094924, and the CS-MACH1 project under Grant Agreement No. 101214613. The ICM-CSIC authors acknowledge the institutional support of the ‘Severo Ochoa Centre of Excellence’ accreditation (CEX2019-000928-S and CEX2024-001494-S funded by AEI 10.13039/501100011033). The opinions expressed herein are those of the authors and do not necessarily reflect those of the projects, the European Union or the European Commission.

## Acknowledgments

We thank all volunteer participants who contributed to BioMARató Catalunya as observers, identifiers, validators or local ambassadors (see the full list here: https://zenodo.org/records/17602383). A single participant may contribute to more than one of these roles, and this multiplicity of contributions is central to the community-based nature of the initiative. We also thank the mobilising organisations that supported participant engagement and data acquisition, including the Federació Catalana d’Activitats Subaquàtiques (FECDAS), Plàncton Diving, Oceánicos, Anèl·lides and Xatrac, as well as the local institutions and organisations that helped facilitate activities along the Catalan coast.

## Competing Interests

The author(s) have no competing interests to declare.

## Notes

### Competing Interest Statement

The authors have declared no competing interest.

